# Ribosomal protein database profiling lends clarity to ribosomal protein evolution and mass distribution

**DOI:** 10.1101/2021.10.25.465821

**Authors:** Wenfa Ng

## Abstract

Existence of theoretical ribosomal protein mass fingerprint as well as utility of ribosomal protein as biomarkers in mass spectrometry microbial identification suggests phylogenetic significance for this class of proteins. To serve the above two functions, facile means of identifying and extracting important attributes of ribosomal proteins from proteome data file of microbial species must be found. Additionally, there is a need to calculate important properties of ribosomal proteins such as molecular weight and nucleotide sequence based on amino acid sequence information from FASTA proteome file. This work sought to support the above endeavour through developing a MATLAB software that extracts the amino acid sequence information of all ribosomal proteins from the FASTA proteome datafile of a microbial species downloaded from UniProt. Built-in functions in MATLAB are subsequently employed to calculate important properties of extracted ribosomal proteins such as number of amino acid residue, molecular weight and nucleotide sequence. All information above are output, as a database, to an Excel file for ease of storage and retrieval. Data available from the analysis of an *Escherichia coli* K-12 proteome revealed that the bacterium possess a total of 59 ribosomal proteins distributed between the large and small ribosome subunits. The ribosomal protein ranges in sequence length from 38 (50S ribosomal protein L36) to 557 (30S ribosomal protein S1). In terms of molecular weight distribution, the profiled ribosomal proteins range in weight from 4364.305 Da (50S ribosomal protein L36) to 61157.66 Da (30S ribosomal protein S1). More important, analysis of the distribution of the molecular weight of different ribosomal proteins in *E. coli* reveals a smooth curve that suggests strong co-evolution of ribosomal protein sequence and mass given the tight constraints that a functional ribosome presents. Finally, cluster analysis reveals a preponderance of small ribosomal proteins compared to larger ones, which remains to be a mystery to evolutionary biologists. Overall, the information encapsulated in the ribosomal protein database should find use in gaining a better appreciation for the molecular weight distribution of ribosomal proteins in a species, as well as delivering information for using ribosomal protein biomarkers in identifying particular microbial species in mass spectrometry microbial identification.

**Subject areas:** biochemistry, microbiology, bioinformatics, biotechnology, molecular biology,

**Graphical abstract:** 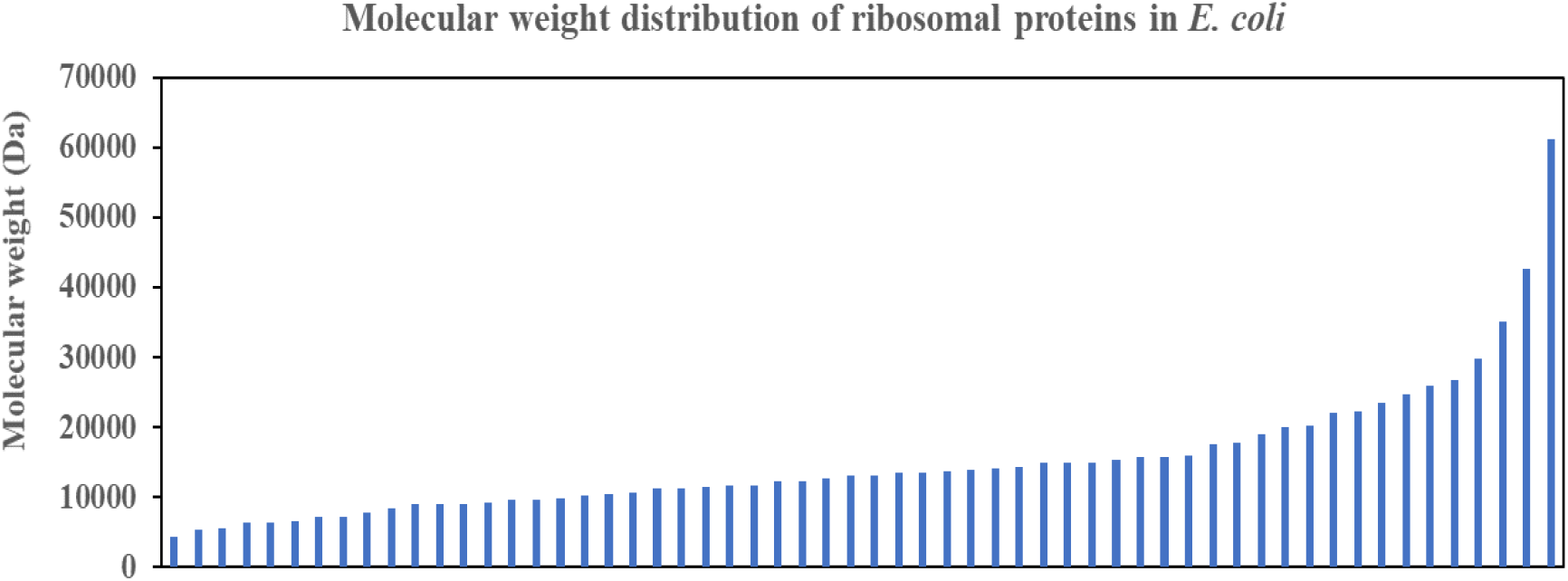

**Short description:** Smooth function describes the distribution of molecular weight of ribosomal proteins in *Escherichia coli*, which suggests strong co-evolution pressure that fine-tunes the molecular weight of individual proteins with the constraint coming from the overall structure of the ribosome that needs to deliver a consistent function to the living cell. Multitude of ribosomal proteins with different roles in the ribosomal proteins under tight co-evolution pressure likely engender the observed smooth curve in the above plot.

Highlights

- Automated MATLAB software for extracting ribosomal protein sequence from FASTA proteome file of a microbial species was developed.
- In-built functions were used to calculate nucleotide sequence, number of residues and molecular weight of each of the extracted ribosomal protein
- All extracted and calculated information were encapsulated as a ribosomal protein database for output to an Excel file for ease of storage and retrieval.
- Information in database could be useful for theoretical ribosomal protein mass fingerprint or help assign ribosomal protein biomarker peaks in matrix-assisted laser desorption/ionization time of flight mass spectrometry (MALDI-TOF MS) microbial identification.

## Introduction

Highly conserved in sequence and function, ribosomal proteins serve important roles in maintaining cellular function through executing their structural roles in constituting the large and small subunit of the ribosomes. Comprising an ensemble of highly defined proteins present in all species in the three domains of life, the family of ribosomal proteins have received much research attention from the perspective of elucidating their individual roles in ribosome biogenesis and function.^1 2^

But, an often less appreciated facet of ribosomal protein biology is their mass distribution. Specifically, recent research examining the possibility of a mass fingerprint traced out by the compendium of ribosomal proteins of a species has garnered a confirmatory “yes” for the existence of ribosomal protein mass fingerprint in all three domains of life.^3^ Such theoretical ribosomal protein mass fingerprints hold implications for their use in the identification of microbial species through mass spectrometry analysis since ribosomal proteins are some of the most highly expressed and abundant proteins in the cellular milieu.^4^ With advances in mass spectrometry instrumentation, ribosomal proteins in whole cell lysate could be analyzed with electrospray ionization (ESI) mass spectrometry^5 6^ or matrix-assisted laser desorption/ionization time of flight (MALDI-TOF) mass spectrometry without an initial protein separation step,^7 8^ which significantly reduces analytical time and increases throughput.

Another aspect in which ribosomal protein could find use in biotechnology is in assisting the identification of microbes profiled through MALDI-TOF MS.^8^ In this usage scenario, the ribosomal protein would serve as biomarkers for identifying specific microbial species whose proteome has been partially interrogated by whole cell MALDI-TOF MS.^4^ Given that ribosomal proteins enjoy relative high abundance, and the mass spectra of microbial whole cells typically comprise high abundance proteins, ribosomal proteins are useful biomarkers for microbial identification purposes.

It is with the above usage scenarios that seed the initial idea for this software development project. Specifically, ribosomal proteins are enmeshed in the thousands of proteins that typified a common proteome file available for a microbial species on UniProt. Automation is certainly necessary to ease the time and burden of extracting the amino acid sequence of all ribosomal proteins of a species for further analysis and processing. To this end, the scripting programming language, MATLAB was employed to develop the necessary code and functions for automated extraction of ribosomal protein amino acid sequence. With sequence in hand, other built-in functions in MATLAB were employed to calculate the nucleotide sequence of the ribosomal protein, number of residues, and molecular weight of protein. The final output is encapsulated as a ribosomal protein database of the species and output to an Excel file for ease of storage and retrieval. As an addon feature, the software also contains a module that would automatically check for wrong residues in the amino acid sequence of the ribosomal proteins. In such cases, no further calculations would be performed on the extracted ribosomal protein amino acid sequence.

The software was applied to the analysis of the *Escherichia coli* K-12 proteome file with the successful extraction of all ribosomal proteins of the species. Analysis of the molecular weight distribution of ribosomal proteins of *E. coli* revealed a smooth curve in mass, which is particularly interesting in that no distinct categories of molecular weight emerged in the analysis. This suggests that the evolution of ribosomal proteins in *E. coli* must have been influenced by the collective ensemble of proteins, with natural selection forces making small adjustments to each protein in order for the overall ribosome to maintain function through cooperative action from different ribosomal proteins in different subunits. Further analysis of distribution of length of each ribosomal protein revealed a preponderance of small ribosomal proteins compared to large ribosomal proteins, which offers clues for further thinking of the evolution of ribosomal proteins in early life on Earth.

## Materials and Methods

The software starts by using fastaread built-in function to read the FASTA format proteome file of the microorganism of interest. Subsequently, the programme cycles through all proteins in the proteome to extract their protein name and amino acid sequence. Prior to calculation of additional properties of the proteins through using MATLAB built-in functions, length (to determine number of residues), molweight (to calculate molecular weight of ribosomal protein), and aa2nt (to calculate nucleotide sequence of ribosomal protein), an error check software was used to check if the extracted amino acid sequence contains letters not belonging to the natural set of amino acids. Those amino acid sequences with extraneous letters would not be used for subsequent calculations. The collated proteome database is subsequently used to profile for ribosomal protein entries, which are extracted and used to build a separate ribosomal protein database. The latter database was subsequently output to an Excel file for ease of storage and retrieval.

## Results and Discussion

As described, the MATLAB software herein delivers two databases to the user: (i) proteome database for all proteins in the proteome, and (ii) a ribosomal protein database. Figure 1 shows the sample output for the ribosomal protein database that comes with protein name, amino acid sequence, number of residues, molecular weight, and nucleotide sequence.

**Figure 1:**
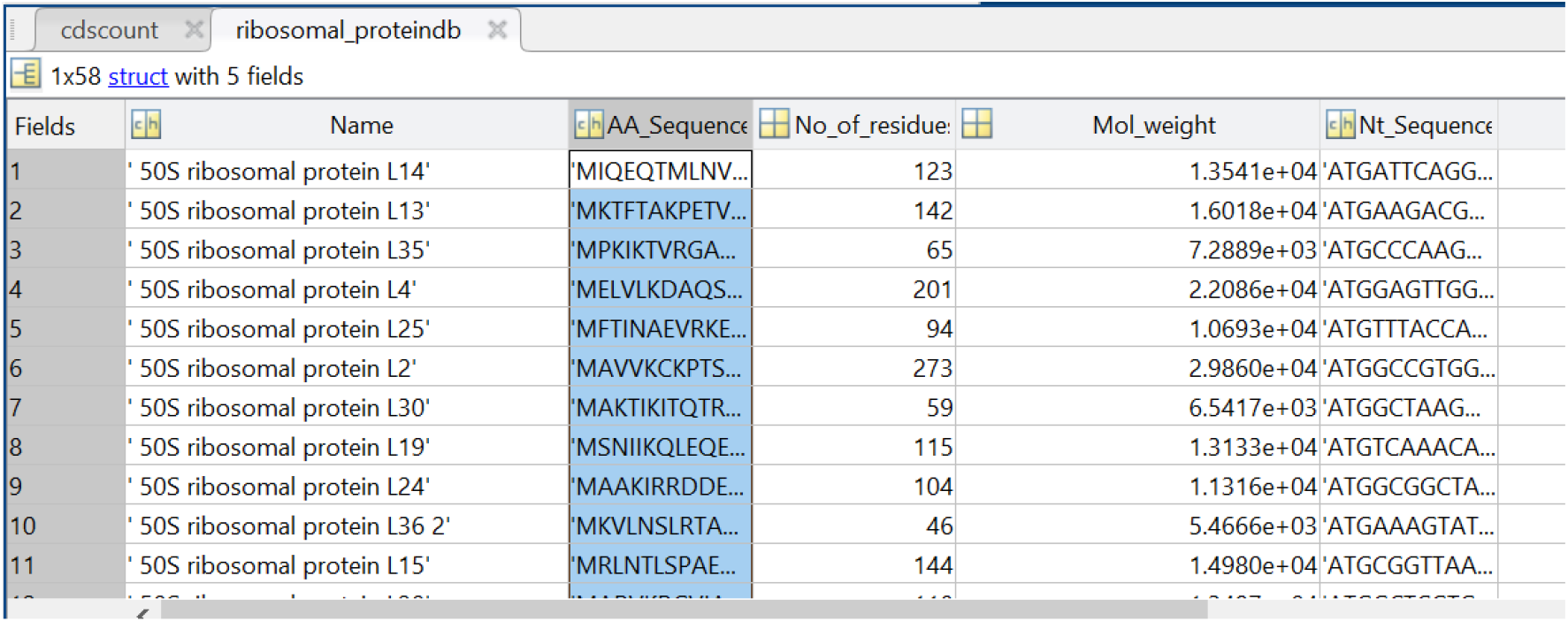
Sample output from the MATLAB software showing the different entries of the ribosomal protein database.

While the ribosomal protein database could be used as a theoretical ribosomal protein mass fingerprint of the bacterium or could inform whether a particular ribosomal protein is useful as biomarker for MALDI-TOF mass spectrometry microbial identification, the data presented in the database is also a rich resource for further analysis that may yield important insights into the biology and evolution of ribosomal proteins.

One approach for analysing the data in the ribosomal protein database of *E. coli* is to understand the distribution of molecular weight of ribosomal proteins across the entire ensemble of proteins. Such analysis would theoretically provide insights into the different categories of mass that different ribosomal proteins fall into, which may help generate further hypothesis into the roles, functions, and evolutionary trajectory of different ribosomal proteins. However, analysis in this direction for *E. coli* throws up an interesting surprise. As shown in Figure 2, the molecular weight distribution of ribosomal proteins in the Gram-negative bacterium does not show distinct mass categories as predicted. Rather, a smooth mathematical function could be used to describe the molecular weight distribution of ribosomal proteins across the entire spectrum of ribosomal proteins profiled.

**Figure 2:**
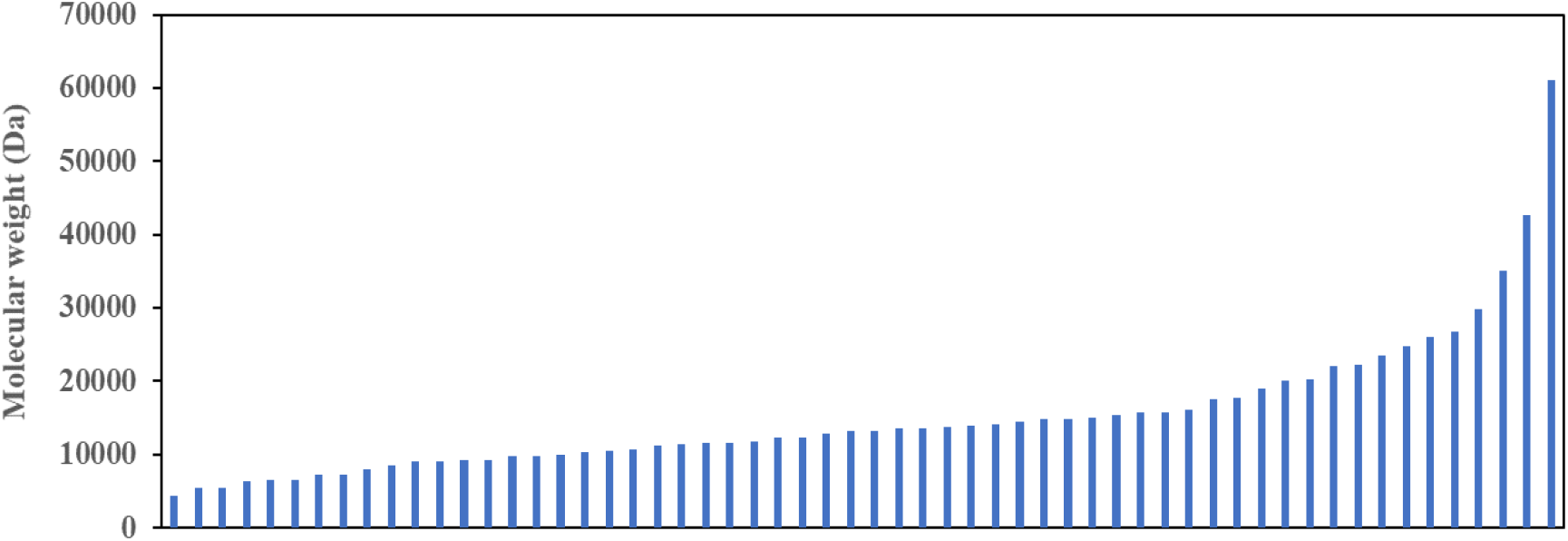
Molecular weight distribution of ribosomal proteins of *Escherichia coli* K-12. Revelation of a smooth curve covering the entire distribution suggests that molecular weight of ribosomal protein was and still is an important selection factor that marks the strong co-evolution of ribosomal proteins with the size and function of the ribosome as major constraining factor.

The results immediate calls into question the idea that ribosomal proteins in a functional ribosome in Bacteria could be binned into distinct categories with possibly different roles and functions for each type of ribosomal protein. Existence of a smooth distribution of molecular weight of ribosomal proteins highlights that molecular weight is an important selection factor during the evolution of ribosomal proteins under the constrains of a size-restricted and function conserved ribosome. Indeed, the obtained molecular weight distribution of ribosomal proteins suggests, on further thinking, strong co-evolution of different ribosomal proteins that occupy distinct locations in the macromolecular complex known as the ribosome. To picture the evolutionary forces pictorially, close proximity between different ribosomal proteins and ribosomal RNA meant that the size of different ribosomal proteins could not deviate much from the optimal settings given the need to maintain function and structure of the ribosome. Such a situation naturally precludes the earlier postulation that ribosomal proteins could be classified into different distinct categories, each with their size limits. If the latter postulation is true, it is hard to fathom how the ribosome macromolecular complex could be constructed with only ribosomal proteins and rRNA, and in its current shape. Overall, the data obtained in Figure 2 dovetails with our conceptual understanding that different ribosomal proteins of different length and size occupy niche, close-interacting positions in the ribosome for ensuring optimal function of this translation machine.

Further analysis of the number of residues and molecular weight of ribosomal proteins through histogram approach (Figure 3) revealed close correspondence between the distribution of ribosomal proteins of different molecular weight and number of residues. This validates the close dependence between number of residues and molecular weight of ribosomal proteins, which marks number of residues as a useful parameter for understanding the relative size of the ribosomal proteins. Predominance of smaller ribosomal proteins in Figure 3a coincides closely with the observation of high abundance of small ribosomal proteins in molecular weight distribution of ribosomal proteins (Figure 2).

**Figure 3:**
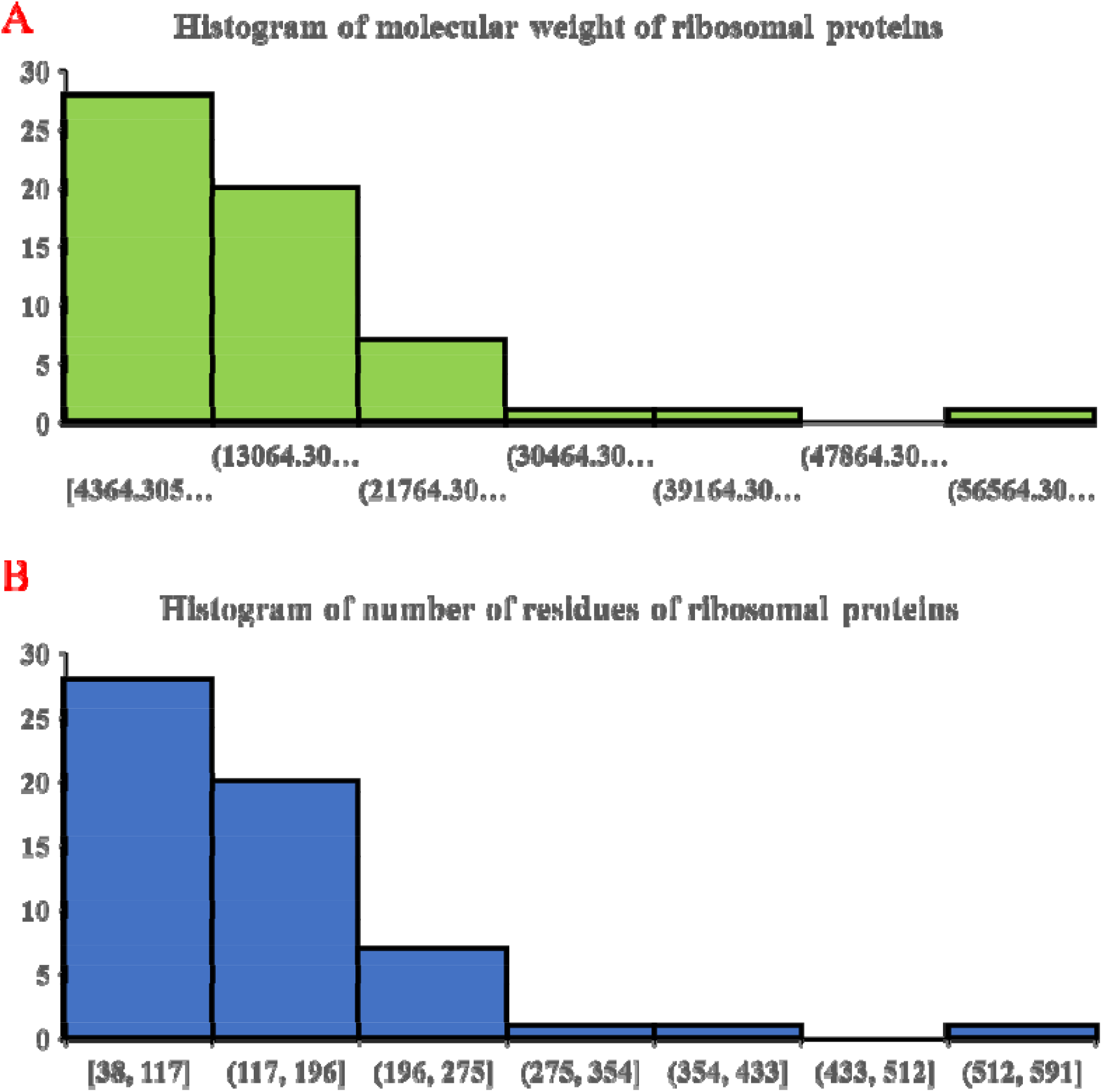
Histogram of molecular weight of ribosomal proteins (A) and number of residues in individual ribosomal proteins (B). Similar histogram for both parameters validate the close correspondence between molecular weight and number of residues in ribosomal proteins. The data also show a preponderance of small ribosomal proteins compared to larger ones.

Overall, the data in Figure 3 highlights the predominance of small ribosomal proteins in the ribosome, which speaks about their utility in lending structure and diversified function to the overall roles of the ribosome. While several large ribosomal proteins could also, in theory, provide the same functions to the ribosome through the properties encapsulated in different protein domains of the same ribosomal proteins, a large intellectual gap in understanding persists in why nature and evolution choose a solution where many small ribosomal proteins divided the task of constituting the ribosome from both the structural and functional perspectives. Given the large number of ribosomal proteins in a ribosome, and presence of multiple protein-protein contact sites in a ribosomal protein, detailed biochemical sleuthing combined with clues from structural biology may be the path forward for unveiling initial aspects of the above problem.

## Conclusions

Molecular weight of ribosomal proteins could find important use as theoretical ribosomal protein mass fingerprint or help identify biomarker peaks in MALDI-TOF mass spectrum of microorganisms. This work aided this endeavour by profiling the relevant information of ribosomal proteins encapsulated in whole proteome of individual microbial species through development of a MATLAB software that extracts and builds a ribosomal protein database. Analysis of the molecular weight and number of residues in ribosomal proteins of *Escherichia coli* using data contained in the ribosomal protein database revealed interesting biological insights that point to strong co-evolution of ribosomal proteins with the confines of the ribosome providing significant evolutionary pressure. Observations of prevalence of small ribosomal proteins in constituting the ribosome macromolecular complex highlights another facet of the evolutionary history of ribosome and its proteins where small ribosomal proteins are chosen over larger ones to lend structure and function to the translation complex.

## Author contribution

The author envisioned the research task, designed and coded the software, performed initial calculations for verifying software performance, and wrote the manuscript.

